# Quantitative real-time PCR assay for the rapid identification of the multidrug-resistant bacterial pathogen *Stenotrophomonas maltophilia*

**DOI:** 10.1101/702985

**Authors:** Tamieka A. Fraser, Mikaela G. Bell, Patrick N.A. Harris, Scott C. Bell, Haakon Bergh, Thuy-Khanh Nguyen, Timothy J. Kidd, Graeme R. Nimmo, Derek S. Sarovich, Erin P. Price

**Author notes:** Corresponding author University of the Sunshine Coast, Locked Bag 4, Maroochydore DC, Qld, 4558, Australia.

## Abstract

*Stenotrophomonas maltophilia* is emerging as an important cause of disease in nosocomial and community-acquired settings, including bloodstream, wound and catheter-associated infections. Cystic fibrosis airways also provide optimal growth conditions for various opportunistic pathogens with high antibiotic tolerance, including *S. maltophilia*. Currently, there is no rapid, cost-effective, and accurate molecular method for detecting this potentially life-threatening pathogen, particularly in polymicrobial specimens, suggesting that its true prevalence may be underestimated. Here, we used large-scale comparative genomics to identify a specific genetic target for *S. maltophilia*, with subsequent development and validation of a real-time PCR assay for its detection. Analysis of 165 *Stenotrophomonas* spp. genomes identified a 4kb region specific to *S. maltophilia*, which was targeted for Black Hole Quencher assay design. Our assay yielded the positive detection of 89 of 89 (100%) clinical *S. maltophilia* strains, and no amplification of 23 non-*S. maltophilia* clinical isolates. *S. maltophilia* was detected in 10/16 CF sputa, demonstrating the utility for direct detection in respiratory specimens. The assay demonstrated good sensitivity, with limits of detection and quantitation on pure culture of ~10 and ~100 genome equivalents, respectively. Our assay provides a highly specific, sensitive, and cost-effective method for the accurate identification of *S. maltophilia*, and will improve the diagnosis and treatment of this under-recognized pathogen by enabling its accurate and rapid detection from polymicrobial clinical and environmental samples.

## Introduction

*Stenotrophomonas maltophilia* is a Gram-negative, intrinsically multidrug-resistant bacterium that is ubiquitous in aqueous environments such as soils, plant roots, water treatments and distribution systems (1). Whilst conventionally overlooked as a laboratory contaminant, or as a common commensal in hospitalized patients, *S. maltophilia* is increasingly being recognized as an important nosocomial pathogen in its own right due to its ability to cause life-threatening disease in immunocompromised individuals (2). This opportunistic pathogen has been isolated from a variety of hospital settings including faucets, sinks, central venous catheters, ice machines, tap water, and water fountains, reinforcing its nosocomial importance (1, 3, 4). *S. maltophilia* most commonly infects people with meningitis, cancer, chronic obstructive pulmonary disease (COPD), or cystic fibrosis (CF), with pneumonia, bacteraemia, and wound and urinary infections being the most frequent clinical manifestations (5, 6). Risk factors for *S. maltophilia* infection include prolonged hospitalization, neutropenia, catheterization, and previous use of broad-spectrum antibiotics (7). The recommended antibiotic treatment for *S. maltophilia* infections is co-trimoxazole; however, resistance towards this antibiotic combination has been documented (2, 8, 9). Indeed, treatment options are limited for *S. maltophilia*, with this pathogen also exhibiting resistance towards several antibiotic classes including fluoroquinolones, macrolides, β-lactams, aminoglycosides, carbapenems, tetracyclines, polymyxins, chloramphenicol and cephalosporins (1, 2). With a mortality rate approaching 70%, the importance of timely identification and effective treatment of *S. maltophilia* infections is paramount (10).

*S. maltophilia* is a common pathogen in CF airways due to its ability to evade many antipseudomonal antibiotics, with chronic *S. maltophilia* infection associated with an increased risk of respiratory disease and mortality (11, 12). CF is an autosomal recessive genetic disorder effecting multiple organs; however, its pathogenesis is most prominent in airways, with ~90% of CF deaths associated with respiratory failure (13). The excessive production of mucus in CF airways provides optimal growth conditions for opportunistic pathogens, which drives most CF morbidity and mortality. Molecular methods have confirmed that CF lower airways harbour diverse microbial communities, with *Pseudomonas aeruginosa* and *Burkholderia cepacia* complex species of greatest concern due to frequent rapid respiratory decline in people infected with these pathogens (14, 15). However, other co-infecting opportunistic pathogens such as *Achromobacter* spp., *S. maltophilia*, *Staphylococcus aureus*, *Haemophilus influenzae* and certain fungal species (e.g. *Aspergillus*) are also prominent in CF airways, and are known to contribute to pathogenesis (14). Indeed, recent studies have shown a mutualistic relationship between *S. maltophilia* and *P. aeruginosa* in CF airways, with compounds produced by *S. maltophilia* under exposure to certain antibiotics, such as imipenem, supporting the survival of otherwise antibiotic-susceptible *P. aeruginosa* strains (16). Furthermore, these *S. maltophilia* compounds can enhance *P. aeruginosa* stress tolerance, increasing polymyxin tolerance (17). These studies highlight the importance of *S. maltophilia* in CF airway pathogenesis, particularly during antibiotic treatment, and emphasize the need for correct species identification in polymicrobial infections.

Although there are a variety of diagnostic methods available for *S. maltophilia* detection, such as PCR amplicon sequencing, VITEK mass spectrometry identification, or key morphological characteristics on growth media, these methods suffer from issues such as limited access to equipment with a large capital expenditure (e.g. ~USD$200,000 for VITEK MS instrumentation), high per-assay cost, the need for highly trained personnel, laboriousness, slow turn-around time, the requirement for purified colonies, or misidentification issues (16, 18, 19). For example, *S. maltophilia* and *P. aeruginosa* exhibit colonial colour differences when grown on bromothymol blue-containing media, which reflects their different metabolic processes (16). However, the use of media containing bromothymol blue is not routine, and thus the retrieval of *S. maltophilia* from polymicrobial specimens requires clinical expertise in identifying appropriate culture media for differentiation of this bacterium from other pathogens. As a non-exhaustive list, *S. maltophilia* has been misidentified as several other organisms, including *Bordetella bronchiseptica*, *Alcaligenes faecalis*, *B. cepacia*, and numerous *Pseudomonas* species, which are common in clinical settings, including CF sputa (1, 20). Diagnostic inconsistencies in *S. maltophilia* detection from clinical specimens can lead to inappropriate or even detrimental treatment (16), particularly for those patients requiring urgent care. There is therefore a need to accurately identify this emerging pathogen to improve antibiotic treatment regimens, stewardship, and patient outcomes. Here, we report the development and validation of a Black Hole Quencher (BHQ) probe-based real-time PCR assay for the specific detection of *S. maltophilia*. Our results indicate that our real-time PCR assay is more sensitive than routine culture for detecting *S. maltophilia*, particularly in polymicrobial respiratory specimens.

## Materials and Methods

### Ethics

This study was approved by The Prince Charles Hospital Human Research Ethics Committee (HREC/13/QPCH/127).

### Genomes of *Stenotrophomonas* spp. and closely related species

All non-redundant *Stenotrophomas* spp. genomes available on the NCBI GenBank database were downloaded as of December 2018. Additional non-redundant *Stenotrophomonas* spp. genomes generated using paired-end Illumina sequencing were downloaded from the NCBI Sequence Read Archive database (SRA; https://www.ncbi.nlm.nih.gov/sra). In total, 165 publicly available genomes were available for this study (Table S1), represented by *S. maltophilia* (including all “*S. pavanii*”; *n*=132), *S. acidaminiphila* (*n*=4), *S. bentonitica* (*n*=3), *S. chelatiphaga* (*n*=1), *S. daejeonensis* (*n*=1), *S. ginsengisoli* (*n*=1), *S. humi* (*n*=1), *S. indicatrix* (*n*=4), *S. koreensis* (*n*=1), *S. lactitubi* (*n*=1), *S. nitritireducens* (*n*=2), *S. panacihumi* (*n*=1), *S. pictorum* (*n*=1), *S. rhizophila* (*n*=2), *S. terrae* (*n*=1), and unassigned *Stenotrophomonas* spp. (*n*=9). Three strains listed on the NCBI database as *Pseudomonas geniculata* (95, AM526, and N1) were included in the phylogenomic analysis to confirm that they were in fact *S. maltophilia*. Sequence data from assembled genomes were converted to simulated 100bp paired-end Illumina reads at 85x coverage using ART version MountRainier (21). SRA data were quality-filtered using Trimmomatic v0.33 (22) using previously described filtering parameters (23) prior to analysis.

### Bioinformatic analysis to identify a *S. maltophilia*-specific genetic locus

The haploid comparative genomics pipeline SPANDx v3.2.1 (24) was used to phylogenetically delineate *S. maltophilia* from non-*S. maltophilia* species, and subsequently, to identify *S. maltophilia*-specific loci. Illumina reads for the publicly available strains (Table S1) were mapped to the closed 4.85Mbp *S. maltophilia* K279a genome (GenBank accession NC_010943.1) (25). Phylogenetic reconstruction of the 165 taxa was performed using 31,246 core, biallelic single-nucleotide polymorphisms (SNPs) using the maximum parsimony function of PAUP* v4.0a.164 (26), rooted with *Stenotrophomonas daejeonensis* JCM 16244 (GenBank accession LDJP01000001.1), and with bootstrapping carried out using 1,000 replicates. To identify genetic loci present in all *S. maltophilia* but absent in all other organisms, including other *Stenotrophomonas* spp., the BEDcov output (27) from SPANDx was examined. Candidate regions were assessed further for specificity and PCR assay design.

### *S. maltophilia* probe-based real-time PCR assay design

Upon identification of a putative *S. maltophilia*-specific genetic target, sequence alignments were used to locate conserved regions. Although some variation was allowed within the amplicon, primer- and probe-binding regions required 100% sequence identity in all *S. maltophilia* strains to avoid false negatives. Oligo self-dimers and heterodimers were assessed *in silico* using NetPrimer (http://www.premierbiosoft.com/netprimer/) and Beacon Designer (http://www.premierbiosoft.com/qOligo/Oligo.jsp/), with configurations resulting in ΔG values of <-8.0 (NetPrimer) and <-4.0 (Beacon Designer) excluded. The following sequences and probe label were chosen for the assay: Smalto-For: 5’-AAGGACAAGGCGATGACCATC, Smalto-Rev: 5’-CCCCACCACGAYTTCATCA, and Smalto-Probe: 5’-FAM-CAGAACGACATCTGGTTGGCG-BHQ1, resulting in an amplicon length of 344 bp. NCBI Microbial Nucleotide discontiguous MegaBLAST (http://blast.ncbi.nlm.nih.gov/) analysis of this amplicon was used to determine assay specificity for only *S. maltophilia*. No mismatches in the probe-binding site were tolerated in any *S. maltophilia* strain, whereas ≥2 mismatches in the probe-binding site were considered sufficient for conferring *S. maltophilia* specificity.

### Microbiological cultures and CF sputum DNA extractions

A total of 89 *S. maltophilia* isolates were obtained for PCR testing: *S. maltophilia* control strain LMG 957, 16 sputum-derived isolates cultured from individuals with CF identified either by amplified rRNA gene restriction analysis (*n*=8 (28)) or *in silico* multilocus sequence typing (*n*=8 (29)), and 72 isolates from various clinical presentations that had previously been identified as *S. maltophilia* by Pathology Queensland Central laboratory, Brisbane, Australia, according to the VITEK2 GNI card (98% of isolates) or VITEK MS (2% of isolates). Strains were grown on Luria-Bertani (LB) agar for 24h at 37 °C. DNA was extracted from a small (toothpick head) swatch of the primary culture by heat soaking in 80 µL of a 5% chelex solution at 95 °C for 10 min, with the resultant DNA diluted 1:10 with molecular grade H_2_O prior to real-time PCR testing.

Sixteen sputa collected from nine Australian adult CF patients underwent DNA extractions using the Zymo Quick-DNA Miniprep Plus Kit (ZYMO Research) protocol, with the addition of an enzymatic lysis solution (20 mg/mL lysozyme, 22U/mL lysostaphin, and 250 U/mL mutanolysin) at 37 °C for 1-3 hours prior to the addition of Proteinase K. Three patients had sputa collected over three time points (range: 13-46 days), and one patient had two sputa collected 320 days apart. “Day 1” samples represent sputa collected on the day of intravenous antibiotic commencement.

### *S. maltophilia* real-time PCR assay validation and specificity testing

Each PCR consisted of 1X Sso Advanced Universal Probes Supermix (Bio-Rad Laboratories, Gladesville, NSW, Australia), optimised primer and probe concentrations of 0.30 and 0.35 µM, respectively (Macrogen Inc., Geumcheon-gu, Seoul, Rep. of Korea), 1 µL of DNA template, and RNase/DNase-free PCR grade water (Thermo Fisher Scientific), to a final per-reaction volume of 5 µL. Thermocycling parameters included an initial hot start activation at 95 °C for 2 minutes, followed by 45 cycles of 95 °C for 3 seconds and 60 °C for 10 seconds, using the CFX96 Touch™ Real-Time PCR Detection System (Bio-Rad). Results were analysed using CFX Maestro v4.1.2433.1219 software.

The 89 *S. maltophilia* isolates and 16 CF sputa were tested using the optimised PCR conditions to ensure accurate detection of *S. maltophilia* DNA in known positive isolates, and in sputum samples from which *S. maltophilia* was variably detected using selective culture (Horse Blood Agar, McConkey Agar, *Burkholderia cepacia* Agar Base and Bacitracin Agar Base). Relative *S. maltophilia* abundance in sputa was determined by performing a previously described 16S rDNA real-time PCR (30) and calculating the cycles-to-threshold difference (∆C_T_) between 16S and the *S. maltophilia* assays. Further specificity testing was performed on 23 non-*S. maltophilia* DNA: *Burkholderia thailandensis* (*n*=1), *Enterobacter aerogenes* (*n*=1), *Enterobacter cloacae* (*n*=4), *Klebsiella oxytoca* (*n*=1), *Klebsiella pneumoniae* (*n*=2), *P. aeruginosa* (*n*=11), *S. aureus* (*n*=1), and *Staphylococcus epidermidis* (*n*=2).

Limit of detection (LoD) and limit of quantification (LoQ) were determined for our newly designed *S. maltophilia* real-time PCR assay using serial dilutions of 50 ng/µL to 0.05 fg/µL *S. maltophilia* DNA across eight replicates/dilution. Twenty-four no-template controls were also included. The upper and lower LoD/LoQ limits were determined as described elsewhere (31).

## Results

### Comparative genomics of *Stenotrophomonas* spp. identifies a *S. maltophilia*-specific gene target

Phylogenomic reconstruction of 165 *Stenotrophomonas* spp. genomes using 31,306 core-genome, biallelic, orthologous SNPs demonstrated a close relationship between *S. maltophilia* and two recently described *Stenotrophomonas* species, *S. indicatrix* and *S. lactitubi* (32), and a clear distinction of these taxa from other *Stenotrophomonas* spp. (Fig 1). This phylogenomic analysis was used to delineate *S. maltophilia* from other species, and to modify incorrect species designations for 43 taxa, including reclassification of all four “*S. pavanii*”, three *P. geniculata*, and 17 *Stenotrophomonas* sp. as *S. maltophilia*, and nine *S. maltophilia* as *Stenotrophomonas* spp. (Table S1).

**Figure 1:**
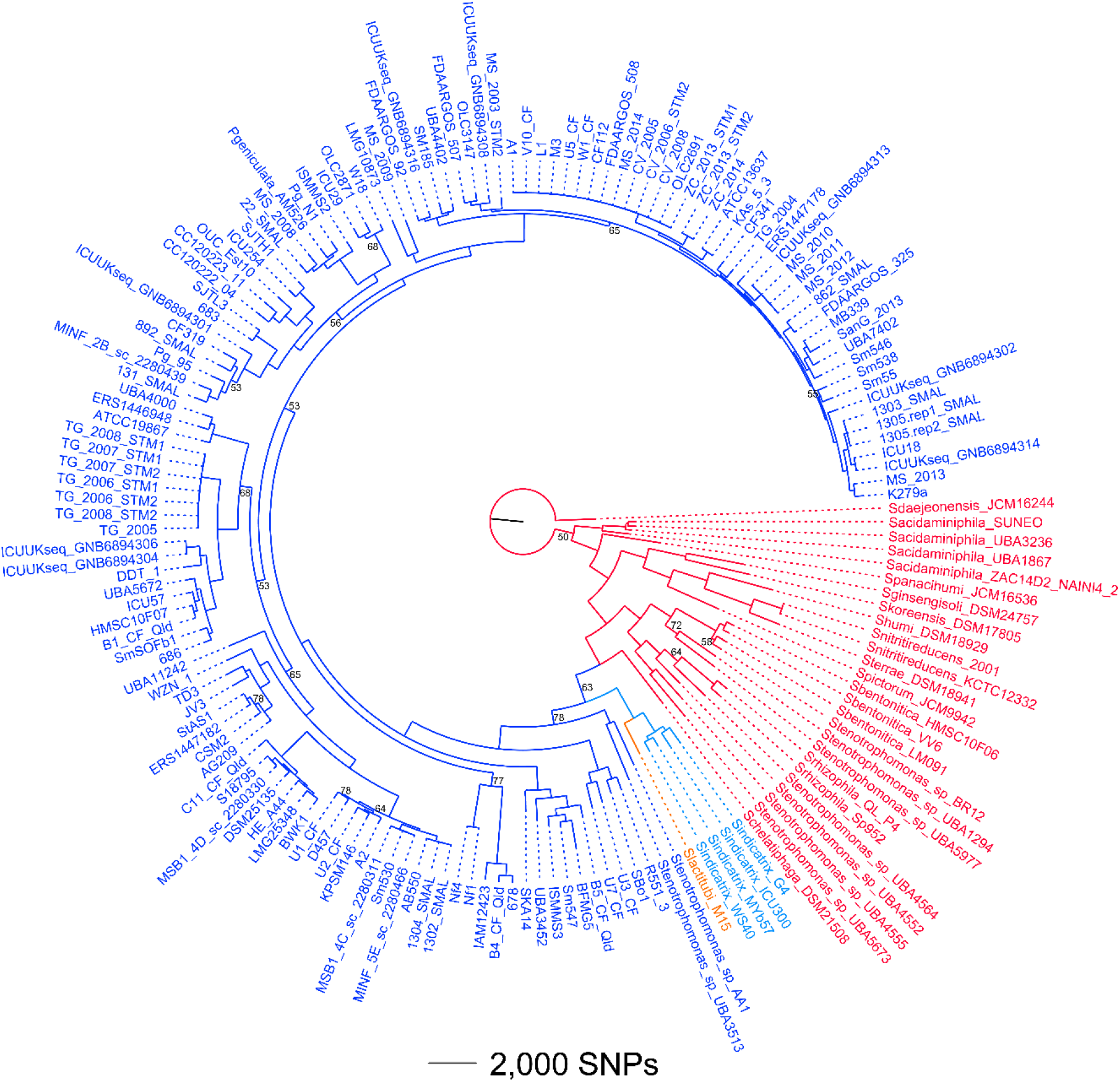
Phylogenomic analysis of *Stenotrophomonas* spp. Dark blue, *S. maltophilia* (target species); red, distantly related *Stenotrophomonas* spp.; light blue, *S. indicatrix*; orange, *S. lactitubi*. The tree was rooted with *S. daejeonensis* JCM16244. Consistency index=0.20. Branches with bootstrap values with <80% support are labelled.

Following identification of the species boundary for *S. maltophilia*, we next identified loci specific for *S. maltophilia*. A single 4kb region was found to be *S. maltophilia*-specific, being present in the 134 *S. maltophilia* genomes but absent or highly divergent in the other *Stenotrophomonas* spp., including *S. indicatrix* and *S. lactitubi*. The genetic coordinates for these loci are 3,935,000-3,939,000 in *S. maltophilia* K279a, spanning gene-coding regions for formate dehydrogenase (α, β, and γ-subunits; encoded by *fdnG, fdnH*, and *fdnI*, respectively).

Upon identification of a suitable 344 bp *S. maltophilia*-specific amplicon, a microbial discontiguous MegaBLAST analysis was conducted to confirm specificity (performed 15Mar19). For all available *S. maltophilia* genomes, 100% sequence identity for both primers and probe was attained. Unexpectedly, the genomes of two recently published *S. indicatrix* strains, which were obtained from soil in Lebanon (RS1 and RS7; GenBank accessions NZ_RKSQ00000000 and NZ_RKSR00000000, respectively), provided close BLAST hits to this amplicon, despite this locus being absent or highly divergent in the four *S. indicatrix* genomes used in the phylogenomic analysis. Fortunately, RS1 and RS7 contained two SNPs in the probe-binding region, including a mismatch at the very 3’ end, which would be expected to inhibit or substantially reduce fluorophore detection in these strains due to poor binding kinetics.

### BLAST analysis identifies further *S. maltophilia* misclassification

In addition to RS1 and RS7, BLAST analysis revealed several non-*S. maltophilia* matches to this amplicon, including three *P. geniculata* strains, several misclassified *Stenotrophomonas* spp. (Table S1), and even misclassifications assigned to distant bacteria e.g. *Acinetobacter baumannii* (4300STDY7045681; GenBank ref. UFGS00000000.1), *Pseudomonas aeruginosa* (E15_London_28_01_14; GenBank ref. CVWF00000000.1) and *Pseudomonas* sp. (UBA10046; GenBank ref. DPXK00000000.1). In all cases, these isolates were confirmed to be *S. maltophilia* according to Microbes BLAST analysis of the whole genome, thereby representing species assignment errors in the NCBI database, and demonstrating the specificity of our *S. maltophilia* target.

### *S. maltophilia* real-time PCR assay specificity

To further assess assay specificity, DNA from 89 *S. maltophilia* isolates derived from both CF and non-CF acute infections was tested. The real-time PCR assay accurately detected 100% of the *S. maltophilia* isolates but did not amplify across any of the tested non-*S. maltophilia* species (*B. thailandensis* [*n*= 1], *P. aeruginosa* (*n*=11), *K. pneumoniae* (*n*=2), *K. oxytoca* (*n*=1), *E. cloacae* (*n*=4), *E. aerogenes* (*n*=1), *S. epidermidis* (*n*=2) and *S. aureus* (*n*=1)). As *S. maltophilia* is known to be a common bacterium in CF sputa, we next tested the assay across 16 CF sputum samples obtained from nine patients. Of these, 10 sputa demonstrated *S. maltophilia* presence at various abundances, with 16S rDNA: *S. maltophilia* ∆C_T_ values ranging from 2.5 (SCHI0019 Day 11; same as pure *S. maltophilia* DNA control) to 17.0 (Table 1). In two longitudinal samples (SCHI0020 and SCHI0021), *S. maltophilia* was only detected at very low levels at the Day 1 time point, with the two subsequent sputa being PCR-negative for this organism. In contrast, *S. maltophilia* persisted in the other two longitudinal samples (SCHI0002 and SCHI0019). There was an 84.6% congruence between the two methods, with two PCR-positive sputa (SCHI0020 Day 1 and SCHI0021 Day 1) being negative by culture (Table 1). However, these two specimens had the lowest *S. maltophilia* load (∆C_T_ values of 13.8 and 17.0), indicating higher sensitivity with the real-time PCR.

**Table 1.** Real-time PCR quantification of *Stenotrophomonas maltophilia* on 16 sputa obtained from cystic fibrosis airways, with concurrent culture diagnosis.

| Sample | $\Delta C_T^{\#}$ | Culture Result |
| --- | --- | --- |
| *S. maltophilia* isolate control | 2.5 | Positive |
| SCHI0002 Day 1 | 7.9 | Positive |
| SCHI0002 Day 320 | 6.0 | Positive |
| SCHI0008 | NA | ND |
| SCHI0010 | NA | Negative |
| SCHI0011 | 12.4 | ND |
| SCHI0013 | 9.7 | ND |
| SCHI0014 | 8.0 | Positive |
| SCHI0019 Day 1 | 6.7 | Positive |
| SCHI0019 Day 11 | 2.5 | Positive |
| SCHI0019 Day 46 | 6.5 | Positive |
| SCHI0020 Day 1 | 17.0 | Negative |
| SCHI0020 Day 6 | NA | Negative |
| SCHI0020 Day 31 | NA | Negative |
| SCHI0021 Day 1 | 13.8 | Negative |
| SCHI0021 Day 5 | NA | Negative |
| SCHI0021 Day 13 | NA | Negative |

<sup>#</sup>Cycles-to-threshold difference; NA, no amplification; ND, not determined.

### *S. maltophilia* real-time PCR assay sensitivity

To assess assay sensitivity, LoD and LoQ values were determined (Figure 2). Using a 10-fold DNA dilution series (50 ng/µL to 0.05 fg/µL), the LoD for this assay is ~5 fg/µL, or ~9 genome equivalents (GEs), and the LoQ is 0.5 pg/µL, or ~94 GEs.

**Figure 2.**
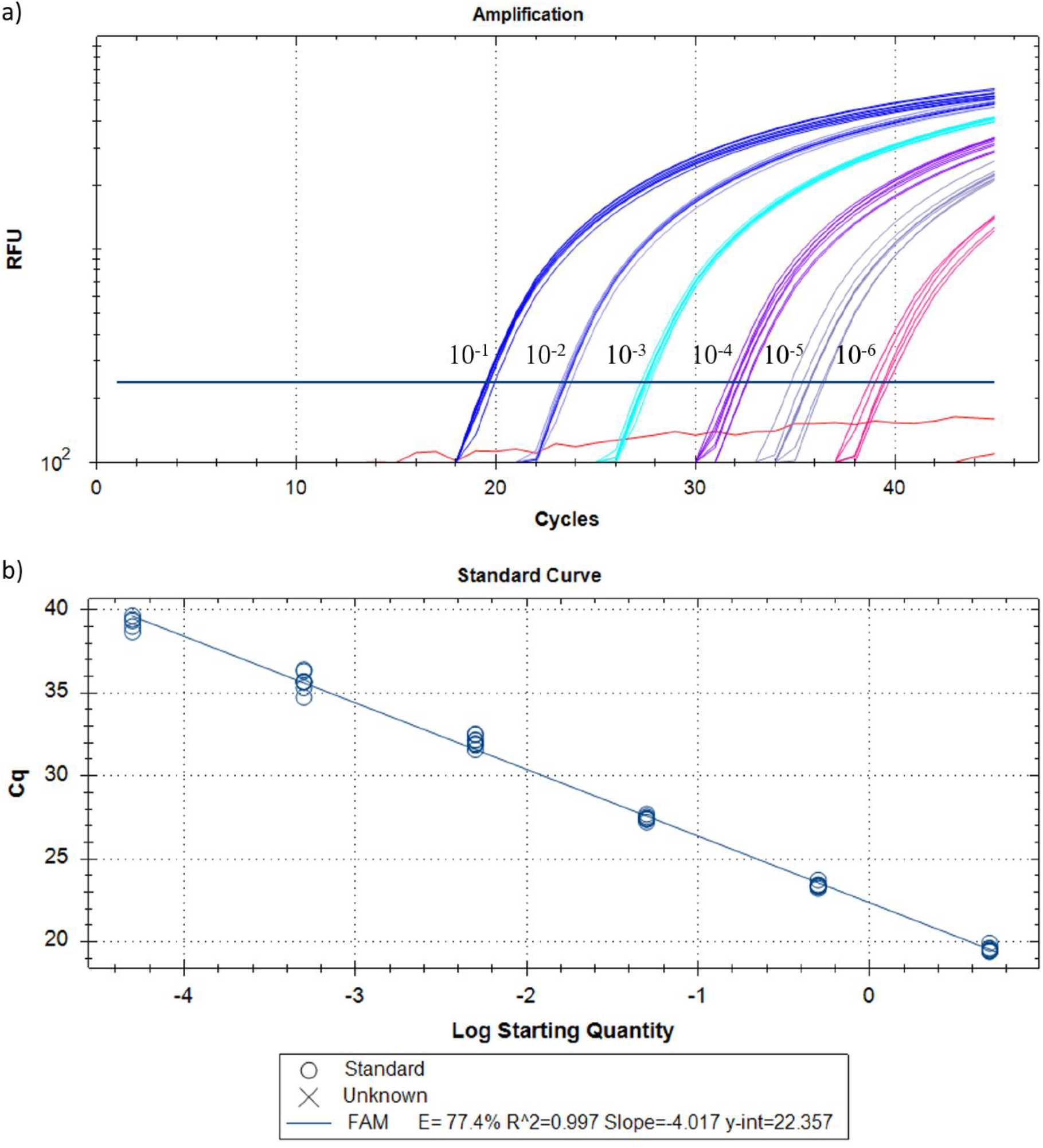
Limits of detection (LoD) and quantitation (LoQ) for the *Stenotrophomonas maltophilia-*specific real-time PCR assay. To determine the LoD and LoQ, serial dilutions of a *S. maltophilia-*positive control DNA sample were performed across eight replicates, ranging from 50 ng/uL to 0.05 fg/µL (a). To assess the correlation coefficient, a standard curve was also included for these dilutions, resulting in an R^2^ of 0.997 (b). LoD and LoQ was identified as ~5 fg/µL and ~0.5 pg/µL, respectively. All 24 negative controls (red) were negative.

## Discussion

*S. maltophilia* is emerging as an important multidrug-resistant nosocomial pathogen, being amongst the top three most common non-fermentative Gram-negative bacilli identified in hospitalized patients (5, 33). Despite *S. maltophilia* being well-adapted to many environments, most infections occur in immunocompromised individuals in the nosocomial setting (10), although community-acquired infections are also on the rise (1). Here, we describe the first real-time PCR assay to detect *S. maltophilia* with 100% accuracy in purified colonies, and demonstrate that this assay is superior to microbiological culture for detecting this multidrug-resistant bacterium in polymicrobial respiratory specimens collected from CF patients.

There is currently a lack of a rapid, cost-effective, accessible, and accurate diagnostic method for *S. maltophilia* detection, particularly from polymicrobial clinical specimens such as CF sputa. As *S. maltophilia* is thought to be the only *Stenotrophomonas* species to cause human disease, mass spectrometry-based systems such as VITEK 2 and VITEK MS are a common diagnostic method in large, centralized pathology laboratories. However, the accuracy of species determination using mass spectrometry is heavily dependent on the quality of the associated databases, and it is currently unknown whether other *Stenotrophomonas* spp. can be accurately differentiated from *S. maltophilia* on these systems. In addition, access to this instrument is limited to well-resourced laboratories owing to a large barrier-to-entry cost (~USD$200,000) (34–36). From a genotyping standpoint, 16S rDNA PCR has been used to identify *S. maltophilia* in blood samples for patients undergoing chemotherapy for leukemia (37), and a multiplex PCR to detect *P. aeruginosa*, *S. maltophilia* and *B. cepacia* successfully identified *S. maltophilia* in 85% of cases (38). However, these assays have either not been optimized to avoid non-specific amplification in other *Stenotrophomonas* spp. and members of the closely related *Xanthomonas* genus, or they require downstream processing (e.g. gel electrophoresis, Sanger sequencing) to confirm results, which is laborious, time-consuming, and raises potential laboratory contamination issues.

Therefore, the purpose of this study was to use large-scale comparative genomics to identify a *S. maltophilia-*specific genetic target, and to subsequently design a highly-specific and accurate real-time PCR-based assay for identifying *S. maltophilia*. Using this approach, we identified a genetic region specific to *S. maltophilia*, which was subsequently targeted for assay development. We found that our newly developed assay correctly identified 89 *S. maltophilia* isolates with 100% accuracy. The accuracy and specificity of this assay is both highly sensitive and selective for *S. maltophilia*, with an LoQ and LoD of ~94 and ~9 GEs, respectively. We chose the Black Hole Quencher probe real-time PCR format due to its relatively inexpensive up-front cost (~USD$25,000 for real-time instrumentation), low per-reaction cost (~USD$0.80 per sample when performed in duplicate), high-throughput capacity, closed-tube format (which eliminates post-PCR contamination concerns), simple set-up, and rapid turn-around-time (~1 h). This format also enables robust identification of target species in polymicrobial specimens. Although not examined in this study, the multi-fluorophore capacity of many real-time PCR instruments also enables multiplexing of probe-based assays for the simultaneous identification of multiple organisms in a single specimen, leading to further cost reductions.

Our *in silico* and laboratory results indicate that all non-*S. maltophilia* microorganisms failed to amplify, with the possible exception of two *S. indicatrix* strains, RS1 and RS7. *S. indicatrix* is a newly identified *Stenotrophomonas* species (32) that has so far been isolated from dirty dishes in Germany (strain WS40; (32)), sewage in China (strain G4; unpublished), a rotting apple in Germany (Myb57; unpublished), soil in Lebanon (strains RS1 and RS7; unpublished), and an unknown source in Germany (ICU300; unpublished). The RS1 and RS7 *S. indicatrix* genomes became available subsequent to assay design; however, BLASTn analysis showed that there were two SNPs in the probe-binding region, including a SNP at the 5’ ultimate base of the BHQ probe, which would likely result in poor or no amplification. Taken together, we show that our assay is highly specific for *S. maltophilia*, particularly in clinical samples, but it also has applicability for testing environmental samples, such as hospital water supplies.

Although the quality of life and life expectancy for people with CF has markedly increased in recent decades due to improvements in antibiotic treatments and clinical management, persistent polymicrobial infections in CF airways remain the primary cause of morbidity and mortality (39). A recent longitudinal study of a single CF patient’s airways using a cutting-edge metatranscriptomic approach, which measures only the ‘active’ microbial population through messenger RNA characterisation, revealed that *S. maltophilia* was the second most prevalent bacterium behind *P. aeruginosa* in the six months prior to death (39). Our results also revealed a high prevalence of *S. maltophilia* in adult CF sputa, with 10/16 samples positive for this bacterium according to our assay. As these sputa samples had concurrent culture results, we demonstrated 84.6% congruence to real-time PCR results, with two culture negative samples returning as positive by real-time PCR (SCHI0020 Day 1 and SCHI0021 Day 1). This finding demonstrates that our assay has a higher sensitivity for detecting *S. maltophilia* in CF clinical specimens than culture methods. Of four longitudinally collected sputa, one patient (SCHI0019) had *S. maltophilia* at all time points (Days 1, 11, and 46; Table 1) despite intravenous antibiotic therapy during this time, and another patient, SCHI0002, was positive for *S. maltophilia* in samples that were collected nearly 12 months apart (Days 1 and 320), indicating either long-term airway persistence or reinfection with this organism. In one sputum sample from SCHI0019 (Day 11), the ∆C_T_ value between the 16S rDNA and *S. maltophilia* PCRs was identical to that of pure *S. maltophilia* culture (Table 1), indicating that *S. maltophilia* had become the dominant, and potentially sole bacterial species, in this specimen. Although outside the scope of this study, this finding demonstrates the potential for *S. maltophilia* to persist and dominate in CF airways following antibiotic-driven microbiome perturbations, which may have implications for rapid re-infection with more formidable pathogens such as *P. aeruginosa*. A further two patients, SCHI0020 and SCHI0021, had *S. maltophilia* at Day 1, but subsequent sampling (up to Day 31 and 13, respectively) were PCR-negative during the intravenous antibiotic treatment phase, indicating successful eradication of *S. maltophilia* in these cases. Future work will entail testing across larger CF sputum panels, including longitudinal samples, to further examine potential mutualistic relationships between *S. maltophilia* and other pathogens such as *P. aeruginosa*, and on assessing assay performance directly on clinical specimens to further reduce sample processing timeframes.

In conclusion, the ability to accurately, rapidly, and cheaply detect *S. maltophilia* is critical for understanding the prevalence of this underappreciated opportunistic pathogen and for reducing its burden of disease. The implementation of this assay in the clinical setting will enable researchers, clinicians and pathologists to more accurately identify this multidrug-resistant bacterium, particularly in isolates that have been ruled out as other multidrug-resistant Gram-negative pathogens, such as *P. aeruginosa* or *Burkholderia* spp. Finally, the correct and rapid identification of *S. maltophilia* will improve antibiotic stewardship measures by enabling more targeted eradication of this pathogen, and in polymicrobial infections such as those commonly found in CF airways, *S. maltophilia* eradication may reduce the prevalence and persistence of more serious pathogens such as *P. aeruginosa*, leading to improved quality of life and lifespans for people with CF.

## Acknowledgements

We wish to thank Kay Ramsay (formerly QIMR-Berghofer; currently University of Otago) for assistance with *S. maltophilia* identification from CF sputum. This study was funded by the University of the Sunshine Coast and Advance Queensland (awards AQRF13016-17RD2 [DSS] and AQIRF0362018 [EPP]). TJK is the recipient of a National Health and Medical Research Council Early Career Fellowship (1088448).

